# Behavioral Relevance Coding in Human Area 46 Precedes Selective Motor Activation to Action Targets

**DOI:** 10.64898/2026.05.18.725862

**Authors:** Simone Del Sorbo, Fausto Caruana, Ivana Sartori, Veronica Pelliccia, Flavia Maria Zauli, Bianca Della Santa, Francesca Talami, Piergiorgio d’Orio, Davide Albertini, Pietro Avanzini, Maria Del Vecchio

**Affiliations:** School of Advanced Studies, University of Camerino, Camerino (MC), Italy; Istituto di Neuroscienze, Consiglio Nazionale delle Ricerche, Parma, Italy; “Claudio Munari” Epilepsy Surgery Centre, ASST GOM Niguarda, Milan, Italy; Department of Biomedical and Clinical Sciences “L. Sacco”, University of Milan, Milan, Italy; Universita’ di Parma, Dipartimento di Medicina e Chirurgia, Parma, Italy

**Keywords:** Action preparation, Goal-directed behavior, Lateral prefrontal cortex, Motor control, Stereo-electroencephalography

## Abstract

The lateral prefrontal cortex (LPFC) actively contributes to the adaptive control of goal-directed behavior. Evidence suggests that the LPFC encodes the behavioral relevance of stimuli, distinguishing action targets from irrelevant objects; however, how this selectivity emerges over time and integrates within large-scale cortical dynamics underlying action preparation remains unclear. We recorded stereo-electroencephalography (sEEG) activity from 43 patients affected by drug-resistant epilepsy while they prepared to grasp an object and compared it with passive observation of the same object. Gamma-band responses were used to characterize cortical responsiveness and to track the spatiotemporal propagation of activity during action preparation. Neural activity first emerged in the occipitotemporal cortex in both experimental conditions and then progressed along two parallel pathways: an intraparietal and a prefrontal one. The intraparietal pathway showed highly similar dynamics during both action preparation and passive observation, suggesting largely intention-independent visuospatial processing. In contrast, the prefrontal pathway exhibited progressively stronger selectivity for behaviorally relevant objects as activity advanced rostrally. Within this pathway, area 46 exhibited sustained responses selectively associated with action-relevant objects, preceding the similarly selective engagement of premotor and motor regions. Overall, our findings identify area 46 as a key node in whole-brain dynamics that orchestrates action preparation by integrating object relevance into executive control signals guiding premotor and motor engagement.

**NEW & NOTEWORTHY:** Filling a critical gap in system-level accounts of LPFC function, whole-brain recordings reveal that area 46 selectively codes object-related information relevant for action and gates motor region activity only when the object represents a true action target.

## INTRODUCTION

Goal-directed behavior relies on the continuous integration of environmental cues, external rules and constraints, and internal factors such as motivation, prior knowledge, or reward expectations. Together, these elements shape how actions are selected and adjusted over time. By weaving these contextual features into a coherent framework, the brain can evaluate the relevance of incoming sensory information relative to goals, allowing behavior to update dynamically as situational demands shift (1, 2). A key region supporting this integrative process in primates is the lateral prefrontal cortex (LPFC), whose role in goal-directed sensory-motor control (3–12) has been widely documented (see 2, 13–16).

A fundamental property of the LPFC is its capacity to encode information that goes beyond the simple categorization of visual stimuli (17–19), progressively incorporating their relevance for behavior. Seminal evidence demonstrated that LPFC activity increases when visual stimuli acquire behavioral meaning through arbitrary stimulus-response associations defined by task rules (20). The link between this behavioral encoding and motor behavior was first highlighted by Simone and collaborators , who showed that LPFC neurons are selectively active during the manipulation of graspable, task-relevant objects. This encoding was then shown to emerge even before action execution: LPFC activity selectively reflects the observation of action-relevant objects during action preparation, prior to movement onset (22). Finally, LPFC responses were found to be strongly modulated by task demands, with significantly greater activity for action targets compared to task-irrelevant objects (23). Together, these findings support a role of the LPFC in gating visual information based on behavioral relevance, thereby linking object representation to the appropriate motor response along the vision-to-action pathway (15, 16). However, the topographical representation of behavioral relevance is not uniform within LPFC. Basile and coworkers identified a caudo–rostral gradient of selectivity within the LPFC, with progressively greater sensitivity to behaviorally relevant objects rostralward (24), an organization that is consistent with a hierarchical transition from perceptual encoding to contextual representation and action planning (2, 25–27). Despite these advances, the functional significance of the prefrontal preference for behaviorally relevant objects remains elusive, mainly because of the paucity of system-level perspectives integrating LPFC computations within the activity of sensorimotor networks. Yet understanding how this selectivity is embedded within whole-brain dynamics during action preparation is essential for clarifying its specific contribution to goal-directed behavior.

To address this gap, we used stereo-electroencephalography (sEEG) in patients affected by drug-resistant epilepsy, leveraging its ability to record intracerebral activity with high spatiotemporal resolution. By asking participants either to passively observe objects or to prepare actions toward them, we characterized the selectivity of LPFC responses and for behaviorally relevant targets. Furthermore, we were able to situate prefrontal activity within large-scale brain networks, also examining how their processing evolves from object perception to action preparation. This system-level perspective moves beyond local descriptions of LPFC functioning, providing insight into how prefrontal object selectivity integrates with distributed circuits to support flexible, goal-directed behavior. By relating LPFC activity to downstream sensorimotor processing, our approach allows us to investigate how contextual information is represented and transformed across networks to control goal-directed behavior.

## MATERIALS AND METHODS

### Participants

SEEG data were collected from 43 patients (16 left-, 20 right-,7 bilateral-implanted, see Supplemental Table S1) affected by drug-resistant focal epilepsy. Intracerebral electrodes were stereotactically implanted in patients undergoing presurgical evaluation for epilepsy at the “Claudio Munari” Epilepsy Surgery Center (ASST GOM Niguarda, Milan, Italy). Electrode placement was determined solely on clinical grounds, based on seizure semiology, scalp EEG recordings, and neuroimaging data. All patients received detailed information about the implantation procedure and sEEG recordings and provided written informed consent. Stereo-EEG recordings were conducted according to standard methodology to localize the brain regions involved in seizure generation and propagation (28). The study was approved by the Ethics Committee of ASST GOM Niguarda (approval ID 939-2.12.2013). Neurological examination was unremarkable in all patients, and none presented with motor or sensory deficits.

### Electrode implantation and anatomical reconstruction

Each electrode (DIXI Medical, Besançon, France) was positioned using stereotactic coordinates and contained multiple leads. After implantation, cone-beam CT scans were acquired and aligned with preoperative T1-weighted MRI scans. We reconstructed the brain and precisely identified the location of each lead. Preoperative MRIs were segmented to extract pial and white matter surfaces, from which the midthickness surface and sulcal patterns were derived. After projecting each recording lead onto the midthickness surface, individual cortical surfaces were resampled and aligned to a standard template (fs-LR-average). For a more detailed description of the reconstruction procedures, refer to Avanzini et al. (29).

### Experimental design

Patients performed a task designed to investigate action preparation in response to cued target objects (*Action condition, AC*, Fig. 1). The task consisted of 60 trials, 20 for each of three objects (a screw, a nail, or a bolt) placed on a workbench and cued in random order. Each trial began with the participant pressing a button box with the right hand, which triggered the illumination of one of three LEDs, each corresponding to one of the objects. The LED illumination, lasting 2 seconds, indicated the target object for the upcoming action. To prevent anticipatory movements during this period, patients were required to keep the button pressed while fixating on the cued object. Following the cue offset, participants executed the reaching and manipulation (i.e., tighten a screw, beat a nail, or screw a bolt by hand). This sequence was then performed by the experimenter while patients passively observed the cued objects from the same viewpoint (*Visual condition, VC*, Fig. 1). This way, we intended to design two conditions in which the patient’s visual experience remained virtually identical, but whose behavioral relevance differed substantially. The experimental design is described also in Del Vecchio et al., where full methodological details are provided (30, 31).

**Figure 1.**
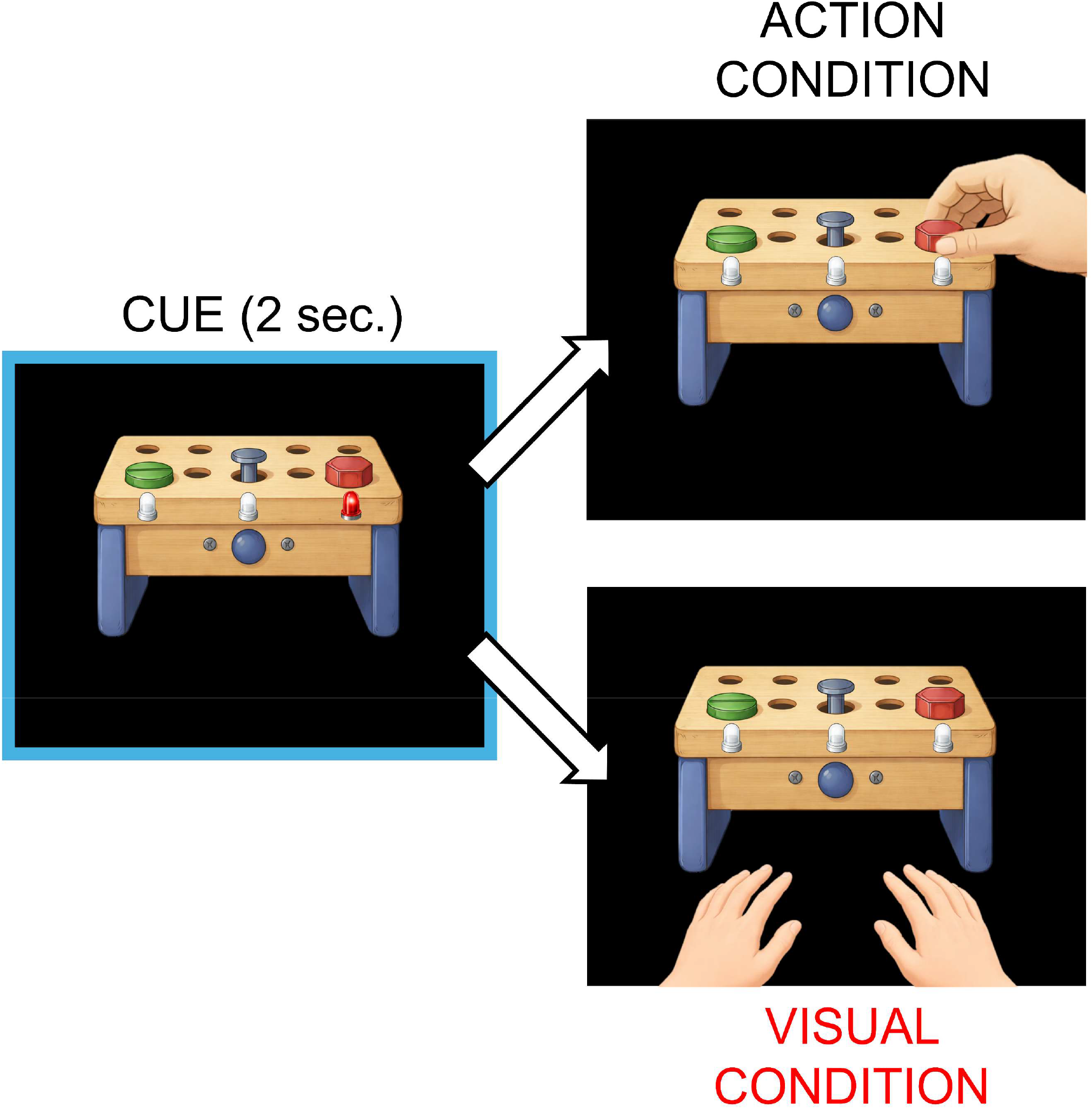
The experimental paradigm. In the AC, each trial began with the participant pressing a button box, triggering a 2-second illumination of one of three LEDs, each cueing a target object (screw, nail, or bolt) on a workbench. Participants maintained the button press without moving until the LED turned off (cue offset), at which point they were free to start the reaching movement followed by the manipulation. In the VC, the same sequence was performed by the experimenter while the participant passively observed the cued objects from the same viewpoint, excluding any motor intention.

### SEEG data recording

For each patient, clinicians selected a neutral intracranial reference based on anatomical and functional criteria. The reference was computed as the average of two adjacent leads in white matter, selected because they showed no responses to standard clinical stimulations (somatosensory, visual, or auditory) and did not elicit sensory or motor effects during electrical stimulation. sEEG signals were recorded with a Neurofax EEG-1100 system (Nihon Kohden) at a 1 kHz sampling rate.

### Data processing

First, following the reconstruction procedure, we selected all leads located in the grey matter, totaling 2,185 and 2,871 leads in the left and right hemispheres, respectively (see Supplemental Figure S2 and Supplemental Table S3). Then, to exclude potentially pathological responses, we removed all leads located within the epileptogenic zone (listed for each patient in Supplemental Table S1). This resulted in the exclusion of 299 (13.68%) and 402 (14%) leads from the left and right hemispheres, respectively (see Supplemental Figure S2). Data from all the remaining leads (1886 and 2469 leads in the left and right hemisphere, respectively) were decomposed into time-frequency representations using a complex Morlet wavelet decomposition (4 cycles, MATLAB function *timefreq* of *EEGLAB* suite). Gamma-band power (GBP) was extracted throughout the two-second object fixation period in both conditions and segmented into 200 bins of 10 ms across 10 adjacent, non-overlapping 10-Hz frequency bands (55–145 Hz), consistent with previous intracranial studies (30, 31). The same analysis was applied to a baseline period, defined as the 350 ms preceding the LED illumination.

### Statistics and reproducibility

To normalize amplitude across patients and leads, GBP values were converted into z-scores relative to the baseline and tested against zero using a two-tailed t-test (p < 0.01, FDR-corrected). To reduce false positives, only leads showing at least four consecutive time bins with significant activity (i.e., 40 ms) were considered responsive. Responsive leads were further classified based on the direction of gamma-band modulation relative to baseline. Leads meeting the criterion exclusively in the negative range were classified as exhibiting gamma-band decreases, whereas those meeting it exclusively in the positive range were classified as gamma-band increases and further included in the analysis. The cortical surface was parcellated according to the atlas by Glasser et al. (32), and signals were aggregated at the regional level. To ensure regional reliability, only regions containing at least two responsive leads and representing at least 10% of the total leads sampled within that region were retained. From this point onward, these regions will be referred to as responsive regions (RRs). When homologous regions were identified bilaterally, their hemispheric similarity was assessed using Pearson correlation. Regions showing a correlation coefficient r ≥ 0.5 were merged into a single region. Having established lead responsivity for both experimental conditions (AC and VC), we next focused on the AC condition to examine whether RRs shared similar temporal dynamics by computing pairwise Pearson correlations between regional average time courses, resulting in a symmetrical correlation matrix. Only correlations reaching statistical significance (p < 0.05) with |r| > 0.3 were further considered. Hierarchical clustering was performed using Euclidean distance to capture differences in response magnitude, with average linkage adopted as a balanced approach between the extremes of single and complete linkage (33). The optimal number of clusters was determined using the Gap Statistic (range: 2–12), which identified a cutoff of 0.76. This threshold was applied to define contiguous blocks of high correlation, producing our final functional units, which included both multi-area clusters and single RRs.

Although response significance was initially assessed independently for AC and VC to identify responsive leads in each condition, the average of GBP time-courses and subsequent comparisons between AC and VC were conducted using only the leads identified as significant in AC. This approach ensures that comparisons are made on the same set of leads, while avoiding the risk of overlooking similar temporal patterns in VC that do not reach significance independently, given that AC is likely a more salient condition due to the active engagement of subjects (34). Clusters identified in AC were also used to group regions in VC, with time courses obtained by averaging responses across AC-responsive leads within each region.

Finally, for each functional unit, onset latency was estimated to order them temporally. Onset was defined as the mean time of the first significant bin within a sequence of four or more consecutive significant bins, computed across all responsive leads within the unit. Variance explained (R^2^) was then used to classify units into High (r >= 0.7, corresponding R^2^ ≥ 0.50) or Low (r < 0.7, corresponding R^2^ < 0.50) correlation groups, reflecting similar versus condition-divergent temporal dynamics. All data processing and statistical analyses were performed using MATLAB R2022b, and visualization of leads on the cortical surface was conducted with Caret software (35).

## RESULTS

### Coverage and Reactivity

Recordings were obtained from 6605 leads in 43 patients, of which 4355 (1886 in the left and 2469 in the right hemisphere) were localized within cortical gray matter and weren’t part of the epileptogenic zone. Among responsive leads (124 left, 145 right), GBP decreases (52 left, 36 right) were mainly observed in primary somatosensory, parieto-opercular, and visual cortices (see Supplemental Figure S4). Notably, decreases in somatosensory areas were observed exclusively in the AC (100%), whereas 28% of decreases in visual cortices were common to both conditions. In line with previous reports, these suppressive responses are interpreted as reflecting sensory habituation to sustained baseline conditions (e.g., prolonged fixation or button press; (36, 37)). Responsive leads showing a GBP increase were grouped into 77 regions (36 left, 41 right). Of these, 26 regions (10 left, 16 right) were retained based on the selection criteria described in the Methods, constituting the core network examined in the following sections. Notably, three regions - premotor eye field (PEF), frontal eye field (FEF), and area 46 - emerged bilaterally and were merged into single regions, as their left- and right-hemisphere average time courses showed a correlation exceeding the threshold defined in the Methods (area 46: r = 0.56; PEF: r = 0.84; FEF: r = 0.57; see Supplemental Figure S5 for their individual time courses). Overall, the identified network encompassed occipitotemporal regions (e.g., LO3, MT complex), intraparietal regions (e.g., MIP, LIPd, IP2, (32)), as well as primary motor, premotor, and prefrontal cortices. In particular, comparison between AC and VC revealed that while occipitotemporal, and intraparietal areas (along with PEF) were engaged in both conditions, the recruitment of anterior areas, specifically primary motor, premotor, and prefrontal cortices was predominantly specific to the AC, with only sparse activation observed during the VC (Fig. 2). The proportion of responsive leads relative to the total recorded per RR is provided in Supplemental Table S6.

**Figure 2.**
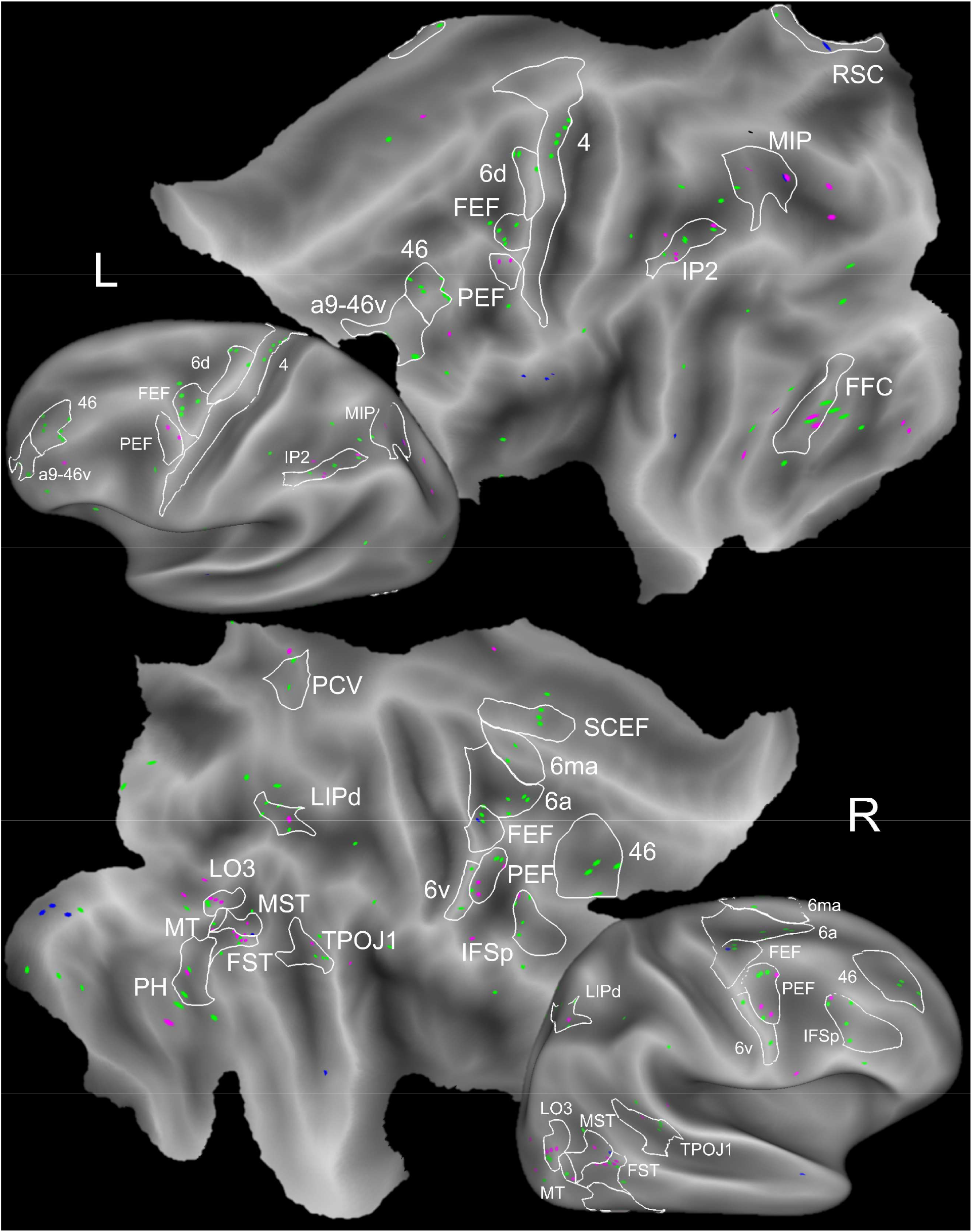
Responsiveness in AC and VC. Responsive leads showing significant GBP increases are shown on flat and inflated cortical maps of the left (L) and right (R) hemispheres for the AC (bright green), VC (blue), and shared between both conditions (magenta). White outlines indicate regions meeting the selection criteria (≥2 responsive leads representing ≥10% of sampled leads in each region) that were included in subsequent analyses.

### Functional network organization and activation dynamics during object observation

To identify coherent functional networks in the AC that might reflect coordinated processing across RRs, we first computed the correlation matrix of the average response profiles for each AC RR and then applied hierarchical clustering. This analysis revealed four distinct clusters (C1–C4), three of which (C1– C3) comprised groups of anatomically contiguous regions (Fig. 3). C1 mainly included occipitotemporal regions (FST, MST, MT, FFC, LO3, PH, TPOJ1, (32)), along with PEF and the supplementary cingulate eye field (SCEF). C2 grouped caudal lateral frontal regions, such as area 6a, FEF, and prefrontal area IFSp, while C3 captured posterior intraparietal regions, including MIP and LIPd. Finally, C4 consisted of the rostral supplementary motor area 6ma and the ventral precuneus (PCV). Individual average time courses of all cluster constituent regions for both conditions are shown in Supplemental Figure S5. Regions that did not cluster with others were treated as independent functional units and analyzed individually, in parallel with the identified clusters. Functional units were then sorted by onset latency (fig. 4A; see Supplemental Table S7). This revealed two parallel activation streams emerging after C1 onset, which exhibited the earliest latency (≅136 ms). One stream progressed caudo-rostrally along the intraparietal cortex, passing through C3 (≅215 ms) and culminating in the anterior intraparietal area IP2 (≅357 ms). The second followed a similar caudo–rostral progression within the prefrontal cortex, starting caudally in C2 (≅201 ms), advancing to area 46 (≅345 ms), and reaching its most anterior segment, area a9/46v (≅630 ms). Within these two streams regions that activated earliest displayed rapid, short-lasting responses, whereas downstream regions in both streams showed more sustained, long-lasting activity (Figure 4C). Activation appeared in C4 at ≅382 ms, preceding that of the retrosplenial cortex (RSC, ≅585 ms). Finally, premotor regions were activated (6d, ≅743 ms; 6v, ≅1270 ms), followed by the primary motor area 4 (≅1798 ms).

**Figure 3.**
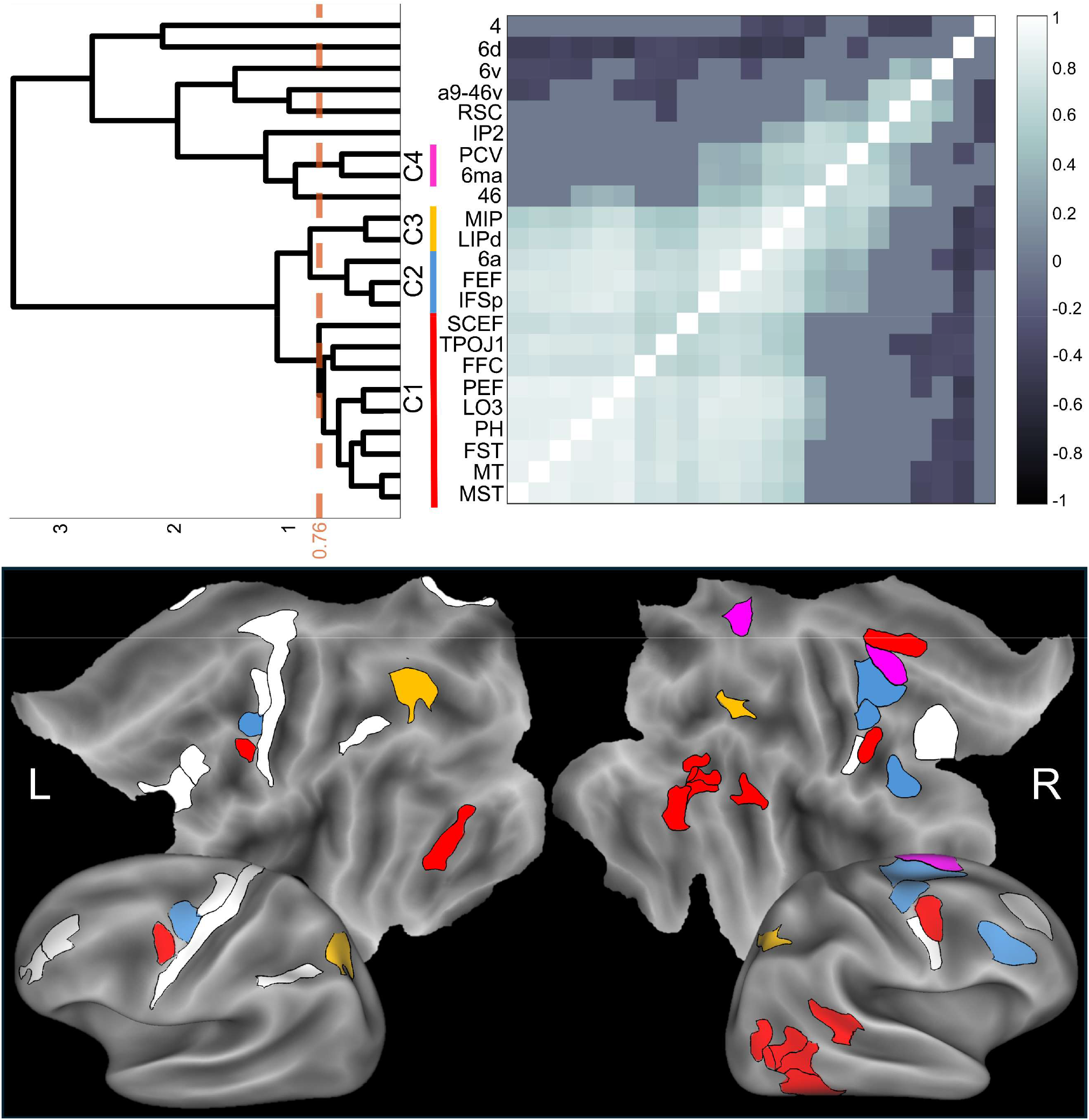
Hierarchical clustering of gamma-band responses in the AC. Top: Hierarchical clustering of the correlation matrix between GBP time courses in the AC. The orange dotted line marks the cutoff threshold (r = 0.76) used to define four clusters (C1–C4), each highlighted in a distinct color. Bottom: Visualization of the four clusters on flat and inflated cortical maps of the left (L) and right (R) hemispheres. Regions are colored according to their cluster assignment. Regions shown in white did not group with any cluster.

**Figure 4.**
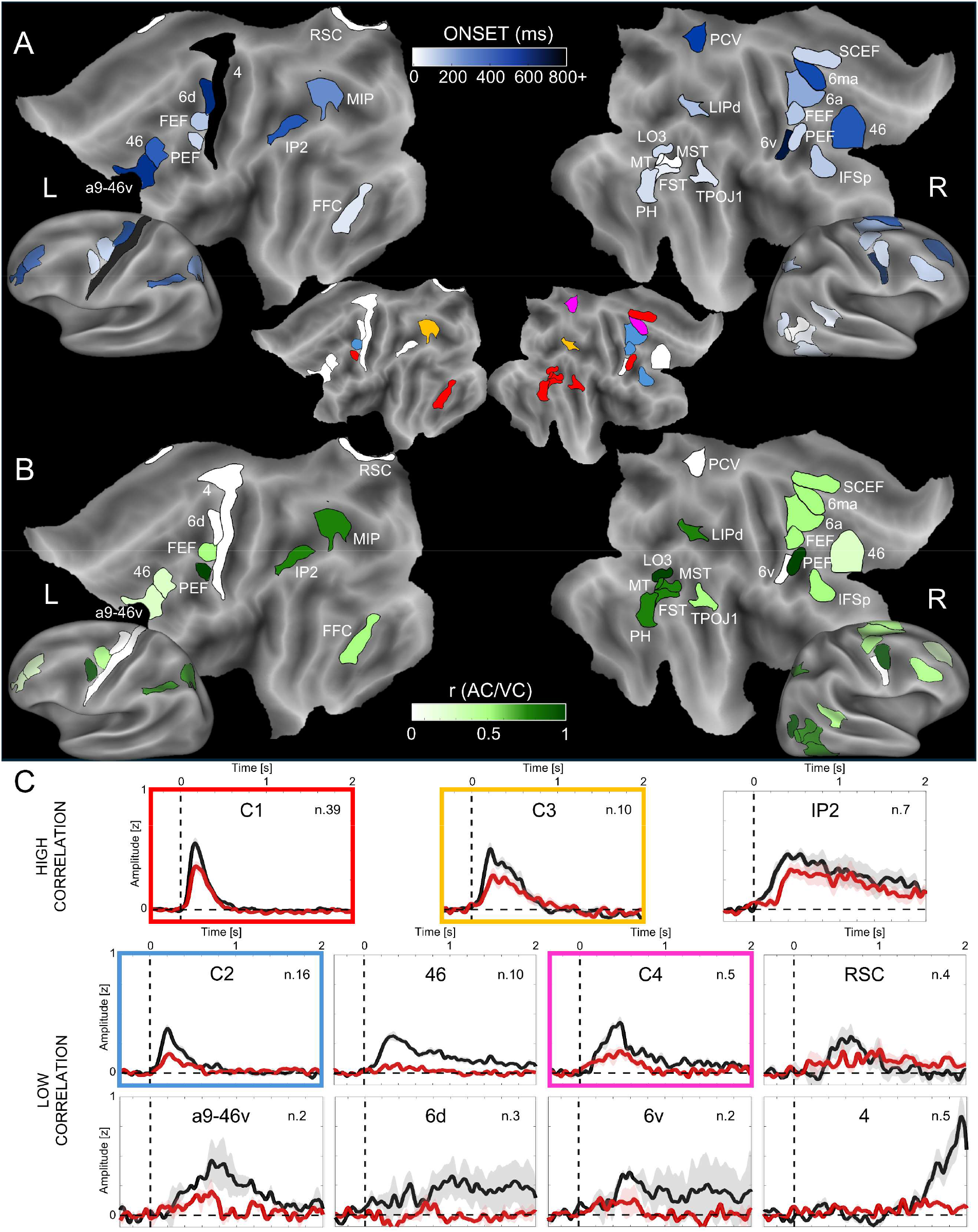
Onset and cross-condition comparison of GBP time courses in AC and VC. (A) Onset latency map. Cortical distribution of onset times for RR in the left (L) and right (R) hemispheres. Onset latency was defined as the mean time point of the first significant bin (within a sequence of at least four consecutive significant bins) across all responsive leads in a given region. Blue shading intensity indicates activation timing, scaling from lighter (earlier) to darker (later) onsets (see scale bar). (B) Cross-condition correlation. Comparison of gamma-band time courses between AC and VC. The intensity of green shading reflects the correlation coefficient (r) between conditions, with darker green indicating stronger similarity (see scale bar). (C) Average time courses (± SE). GBP activity in the AC (black) and VC (red) for all responsive areas. Regions are grouped based on the variance explained (R^2^) between conditions: the top row displays High Correlation group (R^2^ ≥ 0.50, corresponding to r ≥ 0.7), while the bottom rows display Low Correlation group (R^2^ < 0.50, corresponding to r < 0.7). Within each group, panels are ordered by onset latency. Colored frames correspond to the functional clusters, whose cortical maps are reproduced in the center as a visual reference. A moving average filter was applied to the time courses of individual lead within the area (*n* = 5 points).

### Differential response profiles between AC and VC

To compare responses between the AC and VC conditions, we quantified the similarity of their time courses using Pearson correlations for each functional unit (Fig. 4B; see Supplemental Table S7), classifying them into high- and low-similarity groups (Fig. 4C) based on the associated explained variance (see Methods). This analysis revealed a functional dissociation between the two cortical processing streams previously identified. Along the occipitotemporal-intraparietal stream, response profiles were highly similar between AC and VC, indicating that early visual and visuospatial processing operates similarly and largely independently of action intent. In particular, the occipitotemporal cluster (C1) showed strong cross-condition similarity (mean r = 0.72), a pattern that persisted despite differences in the proportion of responsive sites (AC 35.72% vs VC 17.95%). This high similarity persisted along the intraparietal pathway (C3: mean r = 0.79, AC 72.5% vs VC 35%; IP2: r = 0.70, p < .001, AC 63.64% vs VC 27.27%). Even anterior intraparietal areas, despite their later and more sustained activity, responded comparably across conditions, suggesting that these areas transform object information without yet committing to action starting. A contrasting pattern emerged along the prefrontal stream: as information progressed rostrally within the LPFC, activity became increasingly action-selective. Cluster C2 showed moderate similarity (mean r = 0.49) and already favored the AC in responsiveness (AC 23.82% vs VC 2.18%). Selectivity strengthened markedly in area 46 (r = 0.29, p < .001) and in its rostral extension a9/46v (r = 0.32, p < .001), where the complete absence of recruitment during the VC (area 46: AC 28.8% vs VC 0%; area a9/46v: AC 20% vs VC 0%) confirmed that sustained activity in these regions is strictly contingent on the intent to act. A similar preference for the AC was observed in cluster C4 (r = 0.33; AC 17.14% vs VC 0%) and in the retrosplenial cortex (RSC, r = 0.21, p = .003; AC 21.05% vs VC 5.26%). Premotor and primary motor cortices responded almost exclusively during AC (6v: r = 0.09, p = .20, AC 20% vs VC 0%; 6d: r = –0.05, p = .47, AC 25% vs VC 0%; area 4: r = 0.19, p = .008, AC 31.25% vs VC 0%). The near-zero and negative r values observed for 6v and 6d, although not statistically significant, are nevertheless consistent with a condition-dependent difference, given that no VC-related activation was detected at any time point in these leads, suggesting that these premotor regions likely exhibited only peri-baseline fluctuations with no VC-related activation, as shown in the scatterplots in Supplemental Figure S5, which displays significance over time (clustered/non-clustered areas), and cluster activation time points. Together, these results reveal a progressive sharpening of selectivity for the AC from posterior visual regions to the DLPFC and, ultimately, motor cortex.

## DISCUSSION

By recording neural activity while participants prepared to act on or passively observe objects, we identify the lateral prefrontal cortex as a site where object representations are transformed into action-relevant signals during action preparation. While occipitotemporal and intraparietal areas provided early, largely condition-invariant descriptions of the visual scene, LPFC activity progressively differentiated action targets from irrelevant stimuli. This transformation culminated in area 46, whose selectivity emerged before the recruitment of premotor and motor cortices. We suggest that the LPFC does not simply relay visual information forward; instead, it modulates it in light of current goals, generating a relevance-based signal on which downstream motor regions ultimately rely. In this view, action preparation may reflect a hierarchical refinement of object information, with the LPFC contributing to the processing step that links perceptual inputs to emerging goal-directed responses.

### Dissociation between parallel cortical pathways and hierarchical LPFC organization

To position our findings within a broader functional framework, we examined how activity unfolded across cortical regions over time. By ordering functional units by onset latency, we segregated two parallel processing streams following the initial activation of the occipitotemporal cluster C1 during action preparation. One stream traveled along the intraparietal sulcus, progressing from cluster C3 to the anterior intraparietal area IP2, while the other evolved within the LPFC, moving from the caudal lateral frontal cluster C2 toward area 46 and ultimately engaging premotor and motor cortices. Comparisons with passive viewing of the same objects (VC) further highlighted a functional dissociation between these pathways: the intraparietal one exhibited highly similar dynamics across conditions, indicating largely intention-independent visuospatial processing, whereas the prefrontal pathway showed progressively stronger selectivity for behaviorally relevant stimuli as activity advanced rostrally, culminating in sharply action-tuned responses in premotor and motor regions. Additional activations in the retrosplenial cortex (RSC) and cluster C4, which showed enhanced selectivity for the AC, may reflect contributions to future-oriented scene construction (38) and to attentional orienting across object features (39). Across both pathways, responses evolved from brief, short-lasting activity to more sustained downstream activations, suggesting a shift from perceptual encoding toward increasingly higher-order control processes. Early, rapid responses in occipitotemporal cortex likely reflect perceptual object analysis and its distribution toward parietal and frontal regions (40, 41), consistent with the proposed role of inferior temporal regions (TEa/m) in routing object-shape and object-identity information to both pathways (42–45). The gradual extension of prefrontal responses toward more rostral regions, along with the orderly progression of selectivity and onset times, further supports a caudo-rostral anatomo-functional gradient within LPFC (24), here revealed also at the temporal level. Caudal LPFC regions, connected to occipitotemporal cortex (43) and exhibiting rapid responses already modulated by behavioral relevance, might serve as the prefrontal gateway for object information incoming from the ventral visual stream. In contrast, mid-LPFC regions, connected to sensorimotor cortices (46–48) and showing more long-lasting and selective activity might contribute primarily to the selection and planning of context-appropriate actions, as previously proposed (25–27). Finally, activity extended into the most rostral prefrontal territories (a9–46v), associated with higher-order cognitive operations such as rule coding (25, 26) and characterized by strong intra-prefrontal connectivity (46–48). These regions displayed later onsets and prolonged selective responses, consistent with their position at the apex of the LPFC hierarchy.

### Proposed Functional Role of Area 46

Between 300 and 400 ms, sustained activity was observed in parallel in the anterior intraparietal area IP2 and area 46, both of which receive object-related input and project to overlapping premotor regions, effectively linking perceptual and motor processing (40–50). Notably, although anterior intraparietal regions are known to modulate premotor activity based on the visual features of objects (51), they did not exhibit selectivity for behavioral relevance in the present data. This suggests that potential actions may be automatically encoded and made available whenever an object is observed. In contrast, area 46 showed clear selectivity for the AC, possibly reflecting a top-down modulatory mechanism: by encoding an object’s behavioral relevance within the current situational context, LPFC activity may determine whether these latent action representations are transformed into motor preparation, ensuring that motor planning occurs only when the observed object is an action target. This interpretation is further supported by the sequential recruitment observed in the LPFC, followed by selective premotor and motor activations for the AC. Clinical evidence aligns with this view: prefrontal syndromes such as utilization behavior, echopraxia, and anarchic hand syndrome highlight the crucial role of these regions in initiating motor responses only in appropriate contexts, avoiding impulsive, automatic, and non-flexible responses (52–54). Traditionally described as failures to inhibit stimulus-driven actions, such disorders may reflect an alteration in the mechanism that determines whether a movement should be prepared. Dysfunctions of this kind could also underlie the excessive action tendencies observed in conditions like Obsessive-Compulsive Disorder (OCD) and Attention-Deficit/Hyperactivity Disorder (ADHD), both associated with altered DLPFC functioning (55, 56). In this view, abnormal prefrontal control may lead to exaggerated motor preparatory activity even in response to neutral or irrelevant stimuli, as reported in OCD (57).

## DATA AVAILABILITY

The selection of patients has been submitted to a series of stringent precautionary measures. The conditions of our ethics approval do not permit public archiving of individual anonymized raw data. Readers seeking access to the data should contact the corresponding author. Access will be granted to named individuals in accordance with ethical procedures governing the reuse of sensitive data. Specifically, requestors must sign a formal agreement confirming that (a) the user may not use the database for any non-academic purpose (b) the document must be signed by a person with a permanent position at an academic institute or publicly funded research institute. Up to five other researchers affiliated with the same institute for whom the signee is responsible may be named at the end of this document which will allow them to work with this dataset. The user may not distribute the database or portions thereof in any way.

## SUPPLEMENTAL MATERIAL

Supplemental Tables. S1, S3, S6, S7: DOI.

Supplemental Figs. S2, S4, S5: DOI.

## AUTHOR CONTRIBUTIONS

FC and PA conceived and designed research; IS, FMZ, VP, and PGO performed experiments; SDS analyzed data; SDS, MDV and PA interpreted results of experiments; SDS prepared figures; SDS, FC, IS, FMZ, VP, PGO, DA, PA, BDS, FT and MDV drafted manuscript, edited and revised manuscript and approved final version of manuscript.

## REFERENCES

1. Cisek P, Kalaska JF. Neural Mechanisms for Interacting with a World Full of Action Choices. Annu Rev Neurosci 33: 269–298, 2010. doi: 10.1146/annurev.neuro.051508.135409.

2. Koechlin E, Summerfield C. An information theoretical approach to prefrontal executive function. Trends Cogn Sci 11: 229–235, 2007. doi: 10.1016/j.tics.2007.04.005.

3. Arrigoni E, Guidali G, Bolognini N, Pisoni A. Frontal connectivity dynamics encode contextual information during action preparation. Neuroscience: 2025.

4. Gallivan JP, McLean DA, Flanagan JR, Culham JC. Where One Hand Meets the Other: Limb-Specific and Action-Dependent Movement Plans Decoded from Preparatory Signals in Single Human Frontoparietal Brain Areas. J Neurosci 33: 1991–2008, 2013. doi: 10.1523/JNEUROSCI.0541-12.2013.

5. Goldenberg G, Spatt J. The neural basis of tool use. Brain 132: 1645–1655, 2009. doi: 10.1093/brain/awp080.

6. Haaland KY, Harrington DL, Knight RT. Neural representations of skilled movement. Brain 123: 2306–2313, 2000. doi: 10.1093/brain/123.11.2306.

7. Hadland KA, Rushworth MFS, Passingham RE, Jahanshahi M, Rothwell JC. Interference with Performance of a Response Selection Task that has no Working Memory Component: An rTMS Comparison of the Dorsolateral Prefrontal and Medial Frontal Cortex. J Cogn Neurosci 13: 1097–1108, 2001. doi: 10.1162/089892901753294392.

8. Hasan A, Galea JM, Casula EP, Falkai P, Bestmann S, Rothwell JC. Muscle and Timing-specific Functional Connectivity between the Dorsolateral Prefrontal Cortex and the Primary Motor Cortex. J Cogn Neurosci 25: 558–570, 2013. doi: 10.1162/jocn_a_00338.

9. Nee DE. Integrative frontal-parietal dynamics supporting cognitive control. eLife 10: e57244, 2021. doi: 10.7554/eLife.57244.

10. Rowe JB. The Prefrontal Cortex shows Context-specific Changes in Effective Connectivity to Motor or Visual Cortex during the Selection of Action or Colour. Cereb Cortex 15: 85–95, 2004. doi: 10.1093/cercor/bhh111.

11. Shima K, Isoda M, Mushiake H, Tanji J. Categorization of behavioural sequences in the prefrontal cortex. Nature 445: 315–318, 2007. doi: 10.1038/nature05470.

12. Zhou Z, Geng J. Preparatory attentional templates in prefrontal and sensory cortex encode target-associated information. eLife 14: RP104041, 2025. doi: 10.7554/eLife.104041.3.

13. Badre D. Cognitive Control. Annu Rev Psychol 76: 167–195, 2025. doi: 10.1146/annurev-psych-022024-103901.

14. Levy R. The prefrontal cortex: from monkey to man. Brain 147: 794–815, 2024. doi: 10.1093/brain/awad389.

15. Miller EK, Cohen JD. An Integrative Theory of Prefrontal Cortex Function. Annu Rev Neurosci 24: 167–202, 2001. doi: 10.1146/annurev.neuro.24.1.167.

16. Tanji J, Hoshi E. Role of the Lateral Prefrontal Cortex in Executive Behavioral Control. Physiol Rev 88: 37–57, 2008. doi: 10.1152/physrev.00014.2007.

17. Everling S, Tinsley CJ, Gaffan D, Duncan J. Selective representation of task‐relevant objects and locations in the monkey prefrontal cortex. Eur J Neurosci 23: 2197–2214, 2006. doi: 10.1111/j.1460-9568.2006.04736.x.

18. Freedman DJ, Riesenhuber M, Poggio T, Miller EK. Categorical Representation of Visual Stimuli in the Primate Prefrontal Cortex. Science 291: 312–316, 2001. doi: 10.1126/science.291.5502.312.

19. Kusunoki M, Sigala N, Nili H, Gaffan D, Duncan J. Target Detection by Opponent Coding in Monkey Prefrontal Cortex. J Cogn Neurosci 22: 751–760, 2010. doi: 10.1162/jocn.2009.21216.

20. Sakagami M, Niki H. Spatial selectivity of go/no-go neurons in monkey prefrontal cortex. Exp Brain Res 100, 1994. doi: 10.1007/BF00227290.

21. Simone L, Rozzi S, Bimbi M, Fogassi L. Movement‐related activity during goal‐directed hand actions in the monkey ventrolateral prefrontal cortex. Eur J Neurosci 42: 2882–2894, 2015. doi: 10.1111/ejn.13040.

22. Bruni S, Giorgetti V, Bonini L, Fogassi L. Processing and Integration of Contextual Information in Monkey Ventrolateral Prefrontal Neurons during Selection and Execution of Goal-Directed Manipulative Actions. J Neurosci 35: 11877–11890, 2015. doi: 10.1523/JNEUROSCI.1938-15.2015.

23. Rozzi S, Bimbi M, Gravante A, Simone L, Fogassi L. Visual response of ventrolateral prefrontal neurons and their behavior-related modulation. Sci Rep 11: 10118, 2021. doi: 10.1038/s41598-021-89500-0.

24. Basile C, Gerbella M, Gravante A, Lapadula A, Rodà F, Simone L, Fogassi L, Rozzi S. Encoding of visual stimuli and behavioral goals in distinct anatomical areas of monkey ventrolateral prefrontal cortex. PLOS Biol 23: e3003041, 2025. doi: 10.1371/journal.pbio.3003041.

25. Nee DE, D’Esposito M. The hierarchical organization of the lateral prefrontal cortex. eLife 5: e12112, 2016. doi: 10.7554/eLife.12112.

26. Nee DE, D’Esposito M. Causal evidence for lateral prefrontal cortex dynamics supporting cognitive control. eLife 6: e28040, 2017. doi: 10.7554/eLife.28040.

27. Riley MR, Qi X-L, Constantinidis C. Functional specialization of areas along the anterior-posterior axis of the primate prefrontal cortex. Cereb Cortex N Y N 1991 27: 3683–3697, 2017. doi: 10.1093/cercor/bhw190.

28. Cardinale F, Rizzi M, Vignati E, Cossu M, Castana L, d’Orio P, Revay M, Costanza MD, Tassi L, Mai R, Sartori I, Nobili L, Gozzo F, Pelliccia V, Mariani V, Lo Russo G, Francione S. Stereoelectroencephalography: retrospective analysis of 742 procedures in a single centre. Brain 142: 2688–2704, 2019. doi: 10.1093/brain/awz196.

29. Avanzini P, Abdollahi RO, Sartori I, Caruana F, Pelliccia V, Casaceli G, Mai R, Lo Russo G, Rizzolatti G, Orban GA. Four-dimensional maps of the human somatosensory system. Proc Natl Acad Sci 113, 2016. doi: 10.1073/pnas.1601889113.

30. Del Vecchio M, Caruana F, Sartori I, Pelliccia V, Zauli FM, Lo Russo G, Rizzolatti G, Avanzini P. Action execution and action observation elicit mirror responses with the same temporal profile in human SII. Commun Biol 3: 80, 2020. doi: 10.1038/s42003-020-0793-8.

31. Del Vecchio M, Caruana F, Zauli FM, Pelliccia V, Sartori I, d’Orio P, Talami F, Del Sorbo S, Albertini D, Rizzolatti G, Avanzini P. An intracranial insight into (the timing of) the action observation network. NeuroImage 327: 121714, 2026. doi: 10.1016/j.neuroimage.2026.121714.

32. Glasser MF, Coalson TS, Robinson EC, Hacker CD, Harwell J, Yacoub E, Ugurbil K, Andersson J, Beckmann CF, Jenkinson M, Smith SM, Van Essen DC. A multi-modal parcellation of human cerebral cortex. Nature 536: 171–178, 2016. doi: 10.1038/nature18933.

33. Murtagh F, Contreras P. Algorithms for hierarchical clustering: an overview. WIREs Data Min Knowl Discov 2: 86–97, 2012. doi: 10.1002/widm.53.

34. Hardwick RM, Caspers S, Eickhoff SB, Swinnen SP. Neural correlates of action: Comparing meta-analyses of imagery, observation, and execution. Neurosci Biobehav Rev 94: 31–44, 2018. doi: 10.1016/j.neubiorev.2018.08.003.

35. Van Essen DC. Cortical cartography and Caret software. NeuroImage 62: 757–764, 2012. doi: 10.1016/j.neuroimage.2011.10.077.

36. Matsuzaki N, Nagasawa T, Juhász C, Sood S, Asano E. Independent predictors of neuronal adaptation in human primary visual cortex measured with high-gamma activity. NeuroImage 59: 1639–1646, 2012. doi: 10.1016/j.neuroimage.2011.09.014.

37. Onishi H, Oyama M, Soma T, Kubo M, Kirimoto H, Murakami H, Kameyama S. Neuromagnetic activation of primary and secondary somatosensory cortex following tactile-on and tactile-off stimulation. Clin Neurophysiol 121: 588–593, 2010. doi: 10.1016/j.clinph.2009.12.022.

38. Vann SD, Aggleton JP, Maguire EA. What does the retrosplenial cortex do? Nat Rev Neurosci 10: 792–802, 2009. doi: 10.1038/nrn2733.

39. Cavanna AE, Trimble MR. The precuneus: a review of its functional anatomy and behavioural correlates. Brain 129: 564–583, 2006. doi: 10.1093/brain/awl004.

40. Thiebaut De Schotten M, Dell’Acqua F, Valabregue R, Catani M. Monkey to human comparative anatomy of the frontal lobe association tracts. Cortex 48: 82–96, 2012. doi: 10.1016/j.cortex.2011.10.001.

41. Verhagen L, Dijkerman HC, Grol MJ, Toni I. Perceptuo-Motor Interactions during Prehension Movements. J Neurosci 28: 4726–4735, 2008. doi: 10.1523/JNEUROSCI.0057-08.2008.

42. Borra E, Belmalih A, Calzavara R, Gerbella M, Murata A, Rozzi S, Luppino G. Cortical Connections of the Macaque Anterior Intraparietal (AIP) Area. Cereb Cortex 18: 1094–1111, 2008. doi: 10.1093/cercor/bhm146.

43. Gerbella M, Belmalih A, Borra E, Rozzi S, Luppino G. Cortical Connections of the Macaque Caudal Ventrolateral Prefrontal Areas 45A and 45B. Cereb Cortex 20: 141–168, 2010. doi: 10.1093/cercor/bhp087.

44. Orban GA, Zhu Q, Vanduffel W. The transition in the ventral stream from feature to real-world entity representations. Front Psychol 5, 2014. doi: 10.3389/fpsyg.2014.00695.

45. Webster MJ, Bachevalier J, Ungerleider LG. Connections of Inferior Temporal Areas TEO and TE with Parietal and Frontal Cortex in Macaque Monkeys. Cereb Cortex 4: 470–483, 1994. doi: 10.1093/cercor/4.5.470.

46. Borra E, Gerbella M, Rozzi S, Luppino G. Anatomical Evidence for the Involvement of the Macaque Ventrolateral Prefrontal Area 12r in Controlling Goal-Directed Actions. J Neurosci 31: 12351–12363, 2011. doi: 10.1523/JNEUROSCI.1745-11.2011.

47. Borra E, Ferroni CG, Gerbella M, Giorgetti V, Mangiaracina C, Rozzi S, Luppino G. Rostrocaudal Connectional Heterogeneity of the Dorsal Part of the Macaque Prefrontal Area 46. Cereb Cortex 29: 485–504, 2019. doi: 10.1093/cercor/bhx332.

48. Gerbella M, Borra E, Tonelli S, Rozzi S, Luppino G. Connectional Heterogeneity of the Ventral Part of the Macaque Area 46. Cereb Cortex 23: 967–987, 2013. doi: 10.1093/cercor/bhs096.

49. Catani M, Dell’Acqua F, Vergani F, Malik F, Hodge H, Roy P, Valabregue R, Thiebaut De Schotten M. Short frontal lobe connections of the human brain. Cortex 48: 273–291, 2012. doi: 10.1016/j.cortex.2011.12.001.

50. Rojkova K, Volle E, Urbanski M, Humbert F, Dell’Acqua F, Thiebaut De Schotten M. Atlasing the frontal lobe connections and their variability due to age and education: a spherical deconvolution tractography study. Brain Struct Funct 221: 1751–1766, 2016. doi: 10.1007/s00429-015-1001-3.

51. Davare M, Rothwell JC, Lemon RN. Causal Connectivity between the Human Anterior Intraparietal Area and Premotor Cortex during Grasp. Curr Biol 20: 176–181, 2010. doi: 10.1016/j.cub.2009.11.063.

52. De Renzi E, Cavalleri F, Facchini S. Imitation and utilisation behaviour. J Neurol Neurosurg Psychiatry 61: 396–400, 1996. doi: 10.1136/jnnp.61.4.396.

53. Pacherie E. The anarchic hand syndrome and utilization behavior: a window onto agentive self-awareness. Funct Neurol 22: 211–217, 2007.

54. Lhermitte F. ‘UTILIZATION BEHAVIOUR’ AND ITS RELATION TO LESIONS OF THE FRONTAL LOBES. Brain 106: 237–255, 1983. doi: 10.1093/brain/106.2.237.

55. Cortese S, Kelly C, Chabernaud C, Proal E, Di Martino A, Milham MP, Castellanos FX. Toward Systems Neuroscience of ADHD: A Meta-Analysis of 55 fMRI Studies. Am J Psychiatry 169: 1038–1055, 2012. doi: 10.1176/appi.ajp.2012.11101521.

56. Schmidtke K, Schorb A, Winkelmann G, Hohagen F. Cognitive Frontal Lobe Dysfunction in Obsessive-Compulsive Disorder. Biol Psychiatry 43: 666–673, 1998. doi: 10.1016/S0006-3223(97)00355-7.

57. Dayan A, Berger A, Anholt GE. Enhanced action tendencies in obsessive-compulsive disorder: An ERP study. Behav Res Ther 93: 13–21, 2017. doi: 10.1016/j.brat.2017.03.005.

