## Supplemental Materials for "Behavioral Relevance Coding in Human Area 46 Precedes Selective Motor Activation to Action Targets"

| **Subject** | **Gender** | **Age at implant** | **Recorded hemisphere** | **Etiology** | **MRI** | **Epileptic zone** | **Handedness** |
| --- | --- | --- | --- | --- | --- | --- | --- |
| SUB_01 | F | 31 | Left | n.a. | Negative | Left antero-mesial temporal lobe | right |
| SUB_02 | F | 38 | Left | FCD IIb temporo-parietal lobes | FCD of left supramarginal gyrus | Left supramarginal gyrus | right |
| SUB_03 | M | 24 | Left | Gliosis | Negative | Left antero-mesial temporal lobe and insulae | right |
| SUB_04 | M | 46 | Left | n.a. | Negative | Left antero-mesial temporal lobe | right |
| SUB_05 | F | 29 | Right | Gliosis | Negative | Right temporal pole and superior-middle temporal sulcus | right |
| SUB_06 | M | 19 | Right | n.a. | FCD of right mesial parietal lobe | Right mesial parietal lobe | right |
| SUB_07 | F | 39 | Right | FCD IIa of superior frontal gyrus | Negative | Right superior frontal gyrus | right |
| SUB_08 | F | 30 | Left | Gliosis | Negative | Left cingulate gyrus and orbito-frontal cortex | right |
| SUB_09 | M | 19 | Left | Gliosis | Negative | Left middle frontal gyrus and orbito-frontal cortex | right |
| SUB_10 | F | 41 | Left | n.a. | Left posterior temporal ischaemic outcome | Left occipital lobe | right |
| SUB_11 | M | 28 | Right | n.a. | Previous surgery on the right antero-mesial temporal lobe | Right superior temporal gyrus | right |
| SUB_12 | F | 27 | Right | Gliosis | Negative | Right antero-mesial temporal lobe | right |
| SUB_13 | F | 21 | Right | Gliosis + hamartoma in the right temporal lobe | Negative | Right mesial temporal lobe and orbito-frontal cortex | right |
| SUB_14 | F | 36 | Left | n.a. | Negative | Left antero-mesial and superior temporal gyrus | right |
| SUB_15 | F | 38 | Right | n.a. | Negative | Not known | right |
| SUB_16 | F | 24 | Bilateral | FCD IIa of right orbito-frontal cortex | Negative | Right fronto-temporal and insular lobes | right |
| SUB_17 | M | 42 | Right | FCD Ia of right inferior frontal gyrus, orbito-frontal cortex | Negative | Right orbito-frontal cortex | right |
| SUB_18 | F | 39 | Left | n.a. | Negative | Left antero-mesial temporal lobe | right |
| SUB_19 | M | 21 | Right | n.a. | Negative | Right occipital lobe | right |
| SUB_20 | F | 28 | Bilateral | Gliosis | Negative | Not known | right |
| SUB_21 | F | 23 | Right | FCD IIa of post central operculum | FCD of right post central gyrus | Right central and post-central operculi | both-handed |
| SUB_22 | F | 43 | Bilateral | Gliosis | Negative | Right superior and middle frontal gyri | right |
| SUB_23 | M | 37 | Right | Gliosis | Negative | Right mesial frontal lobe | right |
| SUB_24 | M | 32 | Bilateral | n.a. | Bilateral PNH | Bilateral mesial temporal lobe and bilateral PNH | right |
| SUB_25 | F | 38 | Right | Gliosis | Negative | Right temporo-occipital lobes | right |
| SUB_26 | F | 44 | Bilateral | n.a. | Right fronto-central post-surgery outcome | Not Known | right |
| SUB_27 | F | 34 | Right | n.a. | Right PNH | Right PNH and temporal lobe | right |
| SUB_28 | M | 35 | Right | Gliosis | Negative | Right antero-mesial temporal lobe | right |
| SUB_29 | M | 32 | Right | Gliosis | Negative | Right antero-mesial temporal lobe | right |
| SUB_30 | M | 29 | Bilateral | FCD IIa of left cingulate gyrus | Negative | Left fronto-central cingulate gyrus | right |
| SUB_31 | M | 28 | Left | Left HS | Left HS | Left antero-mesial temporal lobe | right |
| SUB_32 | F | 40 | Right | n.a. | Negative | Bilateral temporal and insular lobes | right |
| SUB_33 | F | 39 | Bilateral | FCD Ia of right superior frontal gyrus | Negative | Right superior frontal gyrus | right |
| SUB_34 | M | 40 | Left | Gliosis | Left insular cavernoma surgery outcomes | Left temporo-insular lobes | right |
| SUB_35 | M | 44 | Left | Gliosis | Negative | Left antero-mesial temporal lobe and superior temporal gyrus | right |
| SUB_36 | M | 28 | Left | Gliosis | Negative | Left orbito-frontal cortex | right |
| SUB_37 | M | 37 | Right | Gliosis | Negative | Right temporo-occipital lobes | right |
| SUB_38 | M | 23 | Left | Gliosis | Negative | Left superior temporal gyrus and insulae | right |
| SUB_39 | M | 28 | Right | FCD Ia of superior and middle temporal gyri | Negative | Right antero-mesial temporal lobe | right |
| SUB_40 | F | 40 | Right | FCD Ia of orbito frontal cortex and FCD IIb of temporal pole | Negative | Right orbito frontal cortex and temporal lobe | right |
| SUB_41 | M | 44 | Right | Gliosis | Negative | Right temporal lobe | right |
| SUB_42 | F | 46 | Left | n.a. | Negative | Left antero-mesial temporal lobe | right |
| SUB_43 | M | 40 | Left | Gliosis | Negative | Left parietal and insular lobes | right |

**Supplemental Table S1.** Demographic and clinical data of recorded patients***.***

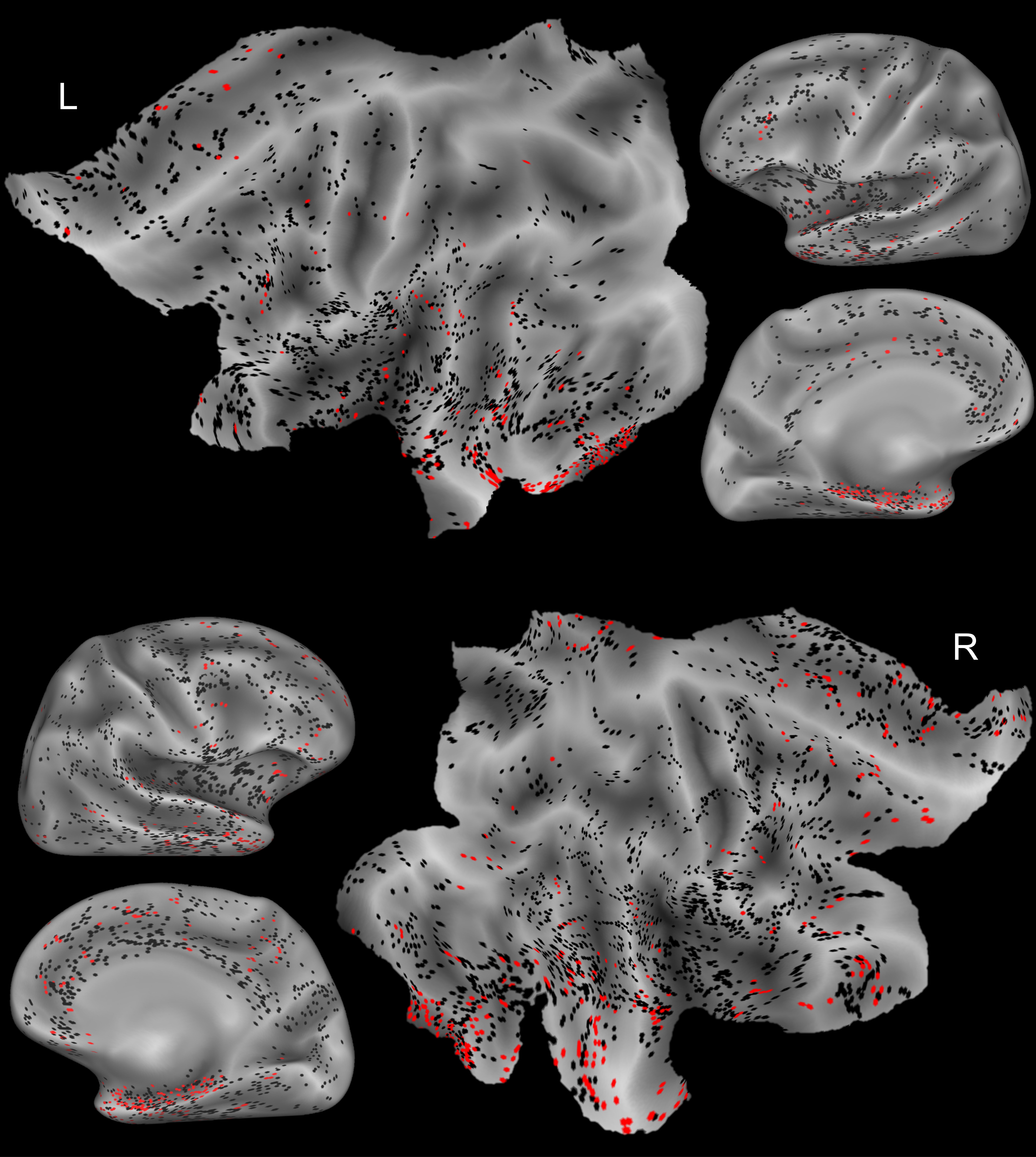

**Supplemental Figure S2.** Cortical localization of recording sites. Sampled sites shown on flat and inflated (lateral and medial) cortical maps of the left (L) and right (R) hemispheres. Leads marked in red were excluded due to their location within or adjacent to epileptogenic zones.

| **Subject** | **# of recording electrodes** | **# of recorded leads** | **# of recorded leads x electrode** | **# of recorded leads in grey matter** | **# of recorded leads in grey matter x electrode** |
| --- | --- | --- | --- | --- | --- |
| SUB_01 | 15 | 148 | 9.6 ± 2 | 124 | 8 ± 2 |
| SUB_02 | 15 | 143 | 9.3 ± 3 | 91 | 5.8 ± 2 |
| SUB_03 | 14 | 172 | 12.1 ± 3 | 133 | 9.3 ± 3 |
| SUB_04 | 16 | 169 | 10.2 ± 2 | 132 | 7.9 ± 2 |
| SUB_05 | 15 | 174 | 11.3 ± 2 | 125 | 8.1 ± 3 |
| SUB_06 | 10 | 143 | 13.9 ± 2 | 102 | 9.8 ± 3 |
| SUB_07 | 16 | 168 | 10.2 ± 3 | 118 | 7.1 ± 3 |
| SUB_08 | 18 | 156 | 8.5 ± 1 | 134 | 7.3 ± 1 |
| SUB_09 | 12 | 130 | 10.8 ± 3 | 101 | 8.4 ± 2 |
| SUB_10 | 16 | 173 | 10.4 ± 2 | 126 | 7.4 ± 2 |
| SUB_11 | 15 | 168 | 11.1 ± 2 | 120 | 7.9 ± 3 |
| SUB_12 | 13 | 170 | 13.0 ± 3 | 131 | 10.0 ± 3 |
| SUB_13 | 19 | 175 | 9.1 ± 2 | 143 | 7.4 ± 3 |
| SUB_14 | 13 | 150 | 11.2 ± 3 | 102 | 7.5 ± 3 |
| SUB_15 | 16 | 159 | 9.8 ± 2 | 139 | 8.6 ± 2 |
| SUB_16 l | 4 | 33 | 8.2 ± 2 | 29 | 7.2 ± 3 |
| SUB_16 r | 16 | 147 | 9 ± 2 | 116 | 7.1 ± 3 |
| SUB_17 | 13 | 137 | 10.5 ± 2 | 124 | 9.5 ± 2 |
| SUB_18 | 15 | 177 | 10.9 ± 3 | 139 | 8.4 ± 2 |
| SUB_19 | 17 | 178 | 10.1 ± 2 | 127 | 7.1 ± 2 |
| SUB_20 l | 10 | 102 | 10.1 ± 3 | 86 | 8.5 ± 2 |
| SUB_20 r | 6 | 55 | 9.2 ± 1 | 50 | 8.3 ± 2 |
| SUB_21 | 14 | 141 | 10.1 ± 2 | 110 | 7.9 ± 2 |
| SUB_22 l | 2 | 27 | 12.5 ± 1 | 13 | 5.5 ± 2 |
| SUB_22 r | 13 | 136 | 10.5 ± 3 | 99 | 7.6 ± 3 |
| SUB_23 | 14 | 161 | 11.4 ± 3 | 122 | 8.6 ± 2 |
| SUB_24 l | 3 | 32 | 10.7 ± 3 | 25 | 8.3 ± 1 |
| SUB_24 r | 15 | 148 | 9.3 ± 3 | 113 | 7.0 ± 2 |
| SUB_25 | 14 | 165 | 11.2 ± 2 | 120 | 8 ± 2 |
| SUB_26 l | 7 | 78 | 10.7 ± 2 | 53 | 7.1 ± 2 |
| SUB_26 r | 7 | 75 | 10.4 ± 2 | 53 | 7.3 ± 2 |
| SUB_27 | 12 | 114 | 9.3 ± 2 | 91 | 7.4 ± 2 |
| SUB_28 | 19 | 190 | 9.9 ± 2 | 131 | 6.8 ± 2 |
| SUB_29 | 13 | 136 | 10.2 ± 3 | 84 | 6.2 ± 2 |
| SUB_30 l | 8 | 80 | 9.8 ± 3 | 59 | 7.1 ± 2 |
| SUB_30 r | 8 | 83 | 10.4 ± 4 | 50 | 6.2 ± 2 |
| SUB_31 | 12 | 126 | 10.2 ± 2 | 95 | 7.7 ± 3 |
| SUB_32 | 12 | 132 | 7.9 ± 2 | 100 | 7.9 ± 2 |
| SUB_33 l | 8 | 79 | 9.2 ± 2 | 68 | 7.9 ± 1 |
| SUB_33 r | 8 | 81 | 10 ± 2 | 60 | 7.4 ± 2 |
| SUB_34 | 16 | 167 | 10.0 ± 2 | 128 | 7.6 ± 3 |
| SUB_35 | 17 | 184 | 9.9 ± 3 | 141 | 7.4 ± 3 |
| SUB_36 | 16 | 165 | 9.9 ± 3 | 133 | 7.9 ± 3 |
| SUB_37 | 13 | 139 | 10.5 ± 2 | 99 | 7.5 ± 3 |
| SUB_38 | 12 | 132 | 10.9 ± 2 | 108 | 8.9 ± 2 |
| SUB_39 | 14 | 156 | 10.9 ± 3 | 99 | 6.8 ± 1 |
| SUB_40 | 15 | 167 | 10.7 ± 3 | 109 | 6.8 ± 2 |
| SUB_41 | 17 | 163 | 9.6 ± 3 | 136 | 8.0 ± 3 |
| SUB_42 | 11 | 121 | 10.6 ± 3 | 74 | 6.4 ± 1 |
| SUB_43 | 13 | 128 | 9.7 ± 2 | 91 | 6.8 ± 2 |

**Supplemental Table S3.** Recording implants details of individual patients.

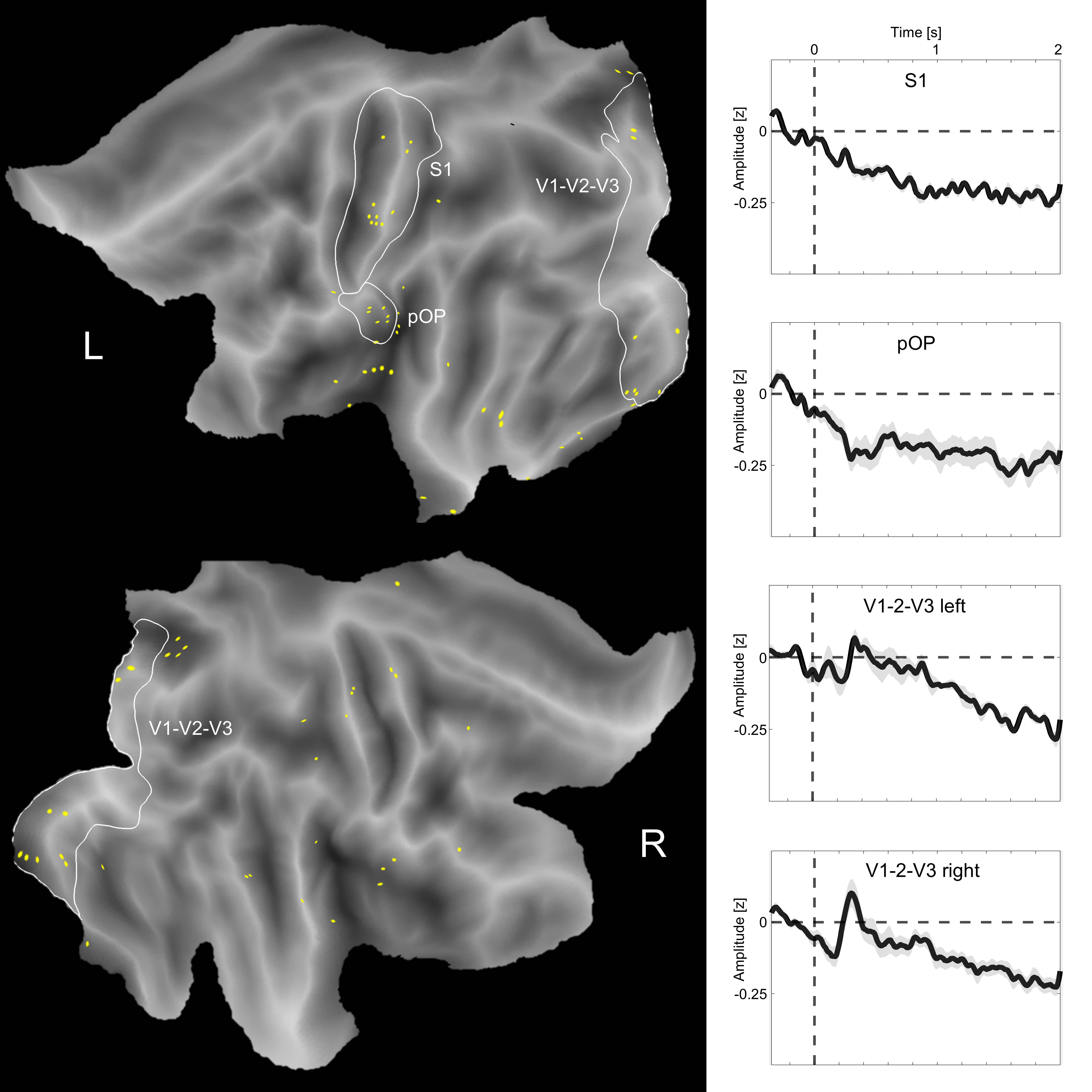

**Supplemental Figure S4.** Leads showing significant GBP decreases. Responsive leads showing significant GBP decreases relative to baseline are displayed in yellow on flat cortical maps of the left (L) and right (R) hemispheres. Average time courses (± SE) for representative areas outlined in white, left primary somatosensory cortex (S1) and parietal operculum (pOP), and bilateral early visual areas (V1-V2-V3), are presented in the panels on the right. A moving average filter was applied to the time courses of individual lead within the area (n= 5 points).

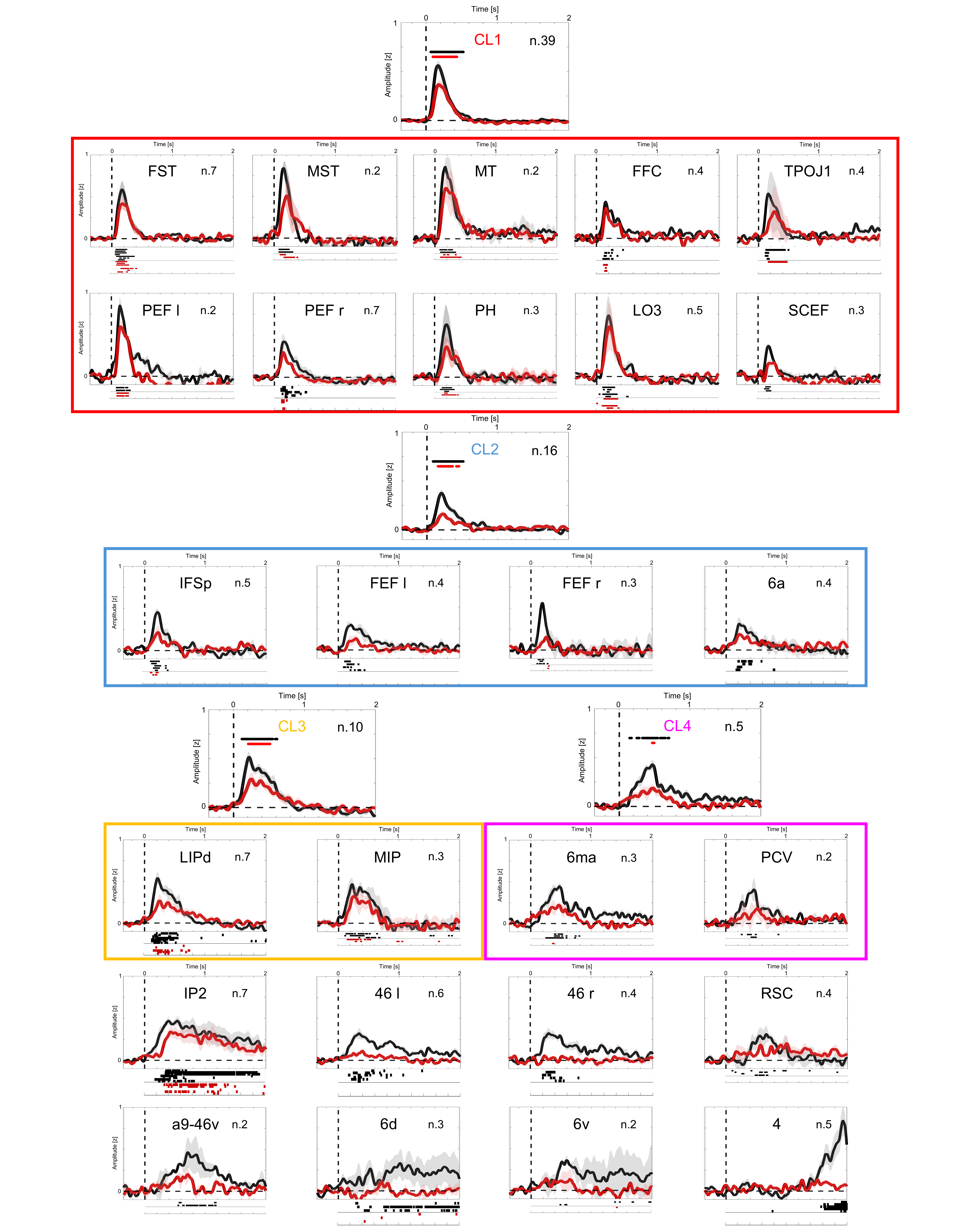

**Supplemental Figure S5.** GBP activity profiles for clusters and individual responsive regions. Average time courses (± SE) of gamma-band activity in the Action (black) and Visual (red) conditions are shown for the four identified clusters (C1–C4) and for all individual responsive areas. Individual regions belonging to specific clusters are grouped within color-coded boxes corresponding to their assigned cluster, while regions that did not cluster with others are displayed at the bottom without framing. Bilaterally responsive areas are displayed separately for each hemisphere. For clusters, asterisks indicate time bins where gamma activity was significantly greater than zero (black asterisks for AC, red for VC; one-tailed t-test, p < 0.01). For individual regions, significant time bins are indicated by dots plotted below each trace (black dots for AC, red dots for VC). A moving average filter was applied to the time courses of individual lead within the area (n= 5 points).

|  | **Left** | | | |  | **Right** | | | |
| --- | --- | --- | --- | --- | --- | --- | --- | --- | --- |
|  | **AC** | | **VC** | |  | **AC** | | **VC** | |
| **Region** | **Responsive / Recorded** | **%** | **Responsive / Recorded** | **%** | **Region** | **Responsive / Recorded** | **%** | **Responsive / Recorded** | **%** |
| **4** | 5/16 | 31.25 | 0/16 | 0 | **46** | 4/18 | 22.22 | 0/18 | 0 |
| **46** | 6/17 | 35.39 | 0/17 | 0 | **6a** | 5/32 | 15.62 | 0/32 | 0 |
| **6d** | 3/12 | 25 | 0/12 | 0 | **6ma** | 3/15 | 20 | 0/15 | 0 |
| **a9-46v** | 2/10 | 20 | 0/10 | 0 | **6v** | 2/10 | 20 | 0/10 | 0 |
| **FEF** | 4/9 | 44.44 | 0/9 | 0 | **FEF** | 3/24 | 12.5 | 1/24 | 4.17 |
| **FFC** | 5/14 | 35.71 | 2/14 | 14.28 | **FST** | 7/21 | 33.33 | 5/21 | 23.81 |
| **IP2** | 7/11 | 63.64 | 3/11 | 27.27 | **IFSp** | 5/22 | 22.73 | 1/22 | 4.55 |
| **MIP** | 3/4 | 75 | 2/4 | 50 | **LIPd** | 7/10 | 70 | 2/10 | 20 |
| **PEF** | 2/4 | 50 | 2/4 | 50 | **LO3** | 5/8 | 62.5 | 3/8 | 37 |
| **RSC** | 4/19 | 21.05 | 1/19 | 5.26 | **MST** | 2/10 | 20 | 3/10 | 30 |
|  | | | | | **MT** | 2/4 | 50 | 1/4 | 25 |
|  |  |  |  |  | **PCV** | 2/14 | 14.28 | 0/14 | 0 |
|  |  |  |  |  | **PEF** | 7/26 | 26.92 | 3/26 | 11.53 |
|  |  |  |  |  | **PH** | 3/10 | 30 | 1/10 | 10 |
|  |  |  |  |  | **SCEF** | 3/13 | 23.07 | 0/13 | 0 |
|  |  |  |  |  | **TPOJ1** | 4/10 | 40 | 1/10 | 10 |

**Supplemental Table S6.** Responsiveness rates across RR. Number of responsive leads showing GBP increases over the total number of recorded leads in each RR for each hemisphere, with corresponding percentages, in the AC and VC. Data are presented separately for the left and right hemispheres.

| **Area** | **n.sub** | **ONSET (ms)** | **r** | **p** |
| --- | --- | --- | --- | --- |
| **MST** | 2 | 90 | 0.76 | <0.001 |
| **MT** | 1 | 95 | 0.81 | <0.001 |
| **FST** | 4 | 116 | 0.84 | <0.001 |
| **PH** | 2 | 143 | 0.71 | <0.001 |
| **LO3** | 2 | 150 | 0.87 | <0.001 |
| **PEF** | 5 | 156 | 0.91 | <0.001 |
| **FFC** | 2 | 127 | 0.53 | <0.001 |
| **TPOJ1** | 2 | 130 | 0.53 | <0.001 |
| **SCEF** | 1 | 157 | 0.50 | <0.001 |
| **IFSp** | 2 | 174 | 0.51 | <0.001 |
| **FEF** | 4 | 179 | 0.50 | <0.001 |
| **6a** | 2 | 225 | 0.46 | <0.001 |
| **MIP** | 3 | 273 | 0.77 | <0.001 |
| **LIPd** | 3 | 190 | 0.80 | <0.001 |
| **46** | 7 | 345 | 0.29 | <0.001 |
| **6ma** | 1 | 377 | 0.47 | <0.001 |
| **PCV** | 2 | 400 | 0.19 | 0.006 |
| **IP2** | 4 | 357 | 0.70 | <0.001 |
| **RSC** | 3 | 585 | 0.21 | 0.003 |
| **a9-46v** | 2 | 630 | 0.32 | <0.001 |
| **6v** | 2 | 1270 | 0.09 | 0.2 |
| **6d** | 2 | 743 | -0.05 | 0.47 |
| **4** | 1 | 1798 | 0.19 | 0.008 |

**Supplemental Table S7.** Regional onset time and response similarity between conditions. For each RR the number of subjects to whom the responsive leads belonged is indicated, and the onset time in AC is reported. The table also includes the Pearson correlation coefficient (r) between GBP regional timecourses in the AC and VC, along with the corresponding significance value of the correlation.
